# p62-DNA-encoding plasmid reverts tumor grade, changes tumor stroma, and enhances anticancer immunity

**DOI:** 10.1101/736686

**Authors:** Franco M. Venanzi, Vladimir Gabai, Francesca Mariotti, Gian Enrico Magi, Cecilia Vullo, Sergey I. Kolesnikov, Alex Shneider

## Abstract

Previously, we reported that the administration of a p62/SQSTM1-encoding plasmid demonstrates high safety and clinical benefits for human cancer patients, having also suppressed tumor growth and metastasis in dogs and mouse models. Here we investigated the mechanistic aspects of these effects. In mammary tumors bearing-dogs, p62 plasmid i.m. injections reduced tumor volumes, and reverted tumor grade to less aggressive lesions in 5 out of 6 animals, with one carcinoma switching to benign adenoma. The treatment increased levels of *alpha*-SMA in stroma cells and collagen 3 in the extracellular matrix, both of which correlate with a good clinical prognosis. p62 treatment also increased the abundance of intratumoral T-cell. To test the role of adaptive immunity, we compared protective effects of the plasmid against B16 melanoma in wild type C57BL/6J mice and in the corresponding SCID strain lacking lymphocytes. The plasmid was only protective in the wild type strain. Also, p62 plasmid amplified anti-tumor effect of adoptive T-cell transfer from tumor-bearing animals to animals challenged with the same tumors. We conclude that the plasmid acts indirectly via re-modeling of the tumor microenvironment, making it more favorable for increased anti-cancer immunity. Thus, the p62-encoding plasmid might be a new adjuvant for cancer treatments.

## Introduction

Recent advances in cancer immunotherapy, particularly immune checkpoint blockade therapy, have dramatically changed the therapeutic strategy against advanced malignancies ^1^. Yet only a subset of patients demonstrate a positive response to any single therapy. Moreover, questions relating to how we can maintain durable clinical responses or how we can successfully treat a broader range of cancers by immunotherapy, remain largely unsolved.

Growing evidence suggests that the major barrier to more successful cancer immuno-/chemotherapy is the tumor microenvironment (TME), where chronic inflammation has a predominant role in tumor survival and proliferation, angiogenesis and immunosuppression ^2–5^. Over the past decades, our understanding of cancer-related inflammation has significantly evolved, and now we have various therapeutic options tailored to the TME ^6^. These therapeutic strategies include inhibiting inflammatory mediators or their downstream signaling molecules, blocking the recruitment of myeloid cells, modulating immunosuppressive functions in myeloid cells and reeducating the TME ^7^. So far, no conclusive studies have been published on stromal content and prognosis in human breast cancer ^8^. In an effort to integrate the effects of the TME and patient outcome into pathological criteria, it has been reported that a “reactive” stromal phenotype may predict breast cancer subtypes with an excellent prognosis ^9^. Indeed, the lowest risk tumors were more likely to have high intratumoral stromal volume-density and high expression of stromal proteins, including alpha-smooth muscle actin (*alpha*-SMA), an actin isoform that marks myofibroblasts and cancer associated fibroblasts. Collagens are critical components of the extracellular matrix regulating tumor progression. Although most research on collagen in breast cancer was focused on type I collagen (Col 1) and its pro-carcinogenic effects ^10 – 12^, new evidence suggest that a related fibrillar type III collagen (Col3) plays an important role in suppressing primary tumor growth and metastasis in a murine model of triple-negative breast cancer ^13, 14^. Even though the role of Col 3 as a co-stimulatory molecules for lymphocytes has not yet been investigated, it’s known that the extracellular matrix may be both a physical barrier to immune cell infiltration and also provide the substratum essential to the interstitial migration of immune cells ^15,16^.

The roles which p62/SQSTM1 plays in cancer and tumor stroma cells remain a subject of active research ^17–20^. A human p62-encoding plasmid was originally proposed as a classic DNA vaccine eliciting adaptive immune response against the p62/SQSTM1 protein, which is reported to be over-expressed in the tumors ^21, 22^. However, the history of science and medicine bears multiple examples when a new phenomenon was explained based on the mechanisms which were most popular at the time they were observed, later turning out to be secondary or insignificant ^23^. Thus, the mechanism of action of the p62 plasmid needs to be reassessed based on the latest observations. Indeed, although treatment with the p62-encoding plasmid was reported as therapeutically beneficial in dogs diagnosed with spontaneous breast cancer ^24^, it turned out later that unlike in humans, most aggressive canine breast tumors show very low or nil p62 expression ^25^.

Therefore, eliciting an anti-p62 adaptive immune response cannot be the only effect of the p62-encoding plasmid. Accordingly, we demonstrate that the p62 DNA treatment dramatically impacted the histopathological features of the malignant lesions. Indeed, after p62 DNA therapy compact tumors appear as multi-lobate neoplasms on an intensive reactive vascular stroma ^24^. The neoplastic lobules were separated and surrounded by thick bands of inflamed fibrous connective, containing scattered aggregates of macrophages, lymphocytes and plasma cells. An increased number of CD3+ intratumoral T-lymphocytes, (CD3+ TIL) was constantly observed ^24^. Taken together, this data raises the question, can a p62 plasmid alter the TME in a way favorable for anti-cancer immune response?

Another line of research revealed that administering the p62 plasmid reduces systemic chronic inflammation in rodent models resulting in the prevention and / or alleviation of numerous inflammation-related diseases. For example, the plasmid reduced metabolic syndrome induced by a high calorie, high fat Western diet ^26^ and an effect on this disease may be linked to anticancer effects ^**27**^. Osteoporosis is also a disorder pertaining to chronic inflammation which shares common signaling pathways with cancer, one example of which being RANK/RANKL signaling ^28^. The p62 plasmid demonstrated both preventive and therapeutic effects in a mouse model of ovariectomy-induced osteoporosis with an effect on pro-inflammatory cytokines and RANK/RANKL signaling ^29^. Importantly, an anti-inflammatory effect of the p62 plasmid may be an indirect effect mediated through a yet unknown third element, as it reported delayed development of age-related macular degeneration (AMD) in rapidly aging OXYS rats ^30^.

Like in humans, dogs’ spontaneous mammary carcinomas are very heterogeneous in terms of morphology and biological behavior ^31^. Dogs with different tumor types and grades demonstrate significant differences in survival. *Simple* and *complex* carcinomas are recorded as the most common type of breast malignancies ^**32**^. Simple carcinoma has a worse prognosis than other mammary tumors, with survival after surgery reduced to 2 years ^33^. Moreover undifferentiated (grade III) simple carcinomas have an increased risk of death when compared with differentiated carcinomas (grade I and II) ^33^. Canine *simple* tumors (e.g. solid and tubulo-papillar subtypes) reveal both histological and molecular homology to human breast carcinomas ^34^. Also, both in canine mammary tumors and in human prostate cancer, the number of intratumoral T-lymphocytes was higher in benign tumors than in their malignant counterparts ^35, 36^. Thus, testing the effect(s) of a cancer treatment on canine model may provide valuable comparative oncology incites.

## Materials & Methods

### p62 DNA Plasmid

Human p62 (Sqstm1, isoform 1) – encoding plasmid was described elsewhere ^41^ and produced endotoxin Free-GMP grade by the Aldevron (ND, USA).

### Dogs Patients and Treatment

Assessment of the therapeutic effect was performed in the veterinary clinic of the University of Camerino (Italy). A total of six dogs, all females, of different breeds and ages were enrolled in the study (Table 1). All of them had histologically confirmed diagnosis of breast carcinoma with WHO stages I-III, progressive disease, and no options for treatment or other treatment options were declined by the owners. The size of tumor was measured every week with a caliper. p62 plasmid was administered i.m. once a weak at the doses of 0.75, 1.5 mg for 3-9 weeks (Table 1). During the treatment, blood was collected for biochemical analysis, and the sizes of tumors were measured manually with calipers according to formula π /6 x L x W x H. and the volume of tumor was calculated. Also, the weight and overall well-being of patients were monitored. All the treatments were performed with full consent of the owners.

**Tab.1.** Patients Characterization. Lesion types: SCa, solid carcinoma; TPca, Tubo-papillary carcinoma

| Pt # | Breed | Age (yrs) | Type | WHO Stage |
| --- | --- | --- | --- | --- |
| 1 | German<br>Seph. | 11 | SCa | T3-N0-M0 |
| 2 | Mongrel | 10 | TP Ca | T1-N0-M0 |
| 3 | German<br>Seph. | 8 | SCa | T3-N2-M1 |
| 4 | Poodle | 14 | SCa | T2-N1-M0 |
| 5 | Breton | 10 | TP Ca | T1-N0-M0 |
| 6 | Boxer | 10 | TP Ca | T2-N0- M0 |

### Tumor Specimens and Immunohistochemistry

Biopsies (Trucut) were performed in all dogs before treatment to establish initial diagnosis. In 5 out of 6 patients, a second biopsies, along with samples from resected tumors (mastectomy) were collected. Patients # 6 had no mastectomy (see the Text). The samples, fixed in 10% neutral buffered formalin, were subjected to histological and immunohistochemical analysis. For each sample 4 μm-thick sections were obtained; one section was stained with haematoxylin-eosin, the other was used for the immunohistochemical analysis. The samples were histologically classified and graded according to criteria of Goldschmidt *et al*. ^32^.

For immunohistochemical analysis sections were mounted on Superfrost®Plus slides and an avidin–biotin–peroxidase-complex (ABC) technique with diaminobenzidine as the chromogen was performed to evaluate the antigen expression. CD3+T cells were stained with rat anti-human CD3 monoclonal antibody (Serotec) and Elite ABC-peroxidase KitsStandard (Vectasain) as previously described ^24^. To further investigate stromal and ECM responses after p62 plasmid administration, slides were stained for alpha-SMA and Col.I and Col III expression by using specific antibodies: Monoclonal Anti-Actin, alpha Smoot Muscle (Sigma). Clone 1A4 (mouse Ig2A isotype): Collagen type I (Novocastra) rabbit polyclonal antibody, and Anti – Col 3A1antibody (Collagen, Type III, alpha 1) Mouse monoclonal (FH-7A) IgG1. All the antibodies were used at the working dilution of 1:75. Sections were counterstained in Mayer’s haematoxylin.

### Mice and Tumor growth assessment

C57BL/6J mice were control (wt) animals, and immunodeficient were -B6. CB17-*Prkdc*^*scid*^/SzJ (C57BL/6 scid). All mice (females 6-8 weeks,18-20 g) were from Jackson Lab. Mice (15 per group) were inoculated with B16 melanoma (3х10^5^ in 0.1 ml of PBS) s.c. in right femurs and injected with p62 plasmid (300 ug/mouse i.m in 0.1 ml of saline), or saline (0.1) as a control on days 1, 8, 15 after tumor inoculation. Our previous experiments demonstrated that empty vector (pcDNA3.1) had no effect on tumor growth as well as saline (FMV unpublished observation). Tumor growth was monitored every other day by a caliper and tumor volume was calculated as indicated above. Statistical analysis was performed by two-way ANOVA with Bonferroni post-tests.

### Preparation of tumor-draining lymph nodes (TDLN) T cells for adoptive immunotherapy

C57BL/6J mice were inoculated subcutaneously with 1 × 10^6^ MCA205 fibrosarcoma cells in both flanks. Twelve days later, inguinal TDLNs were harvested, and single-cell suspensions were prepared and culture-activated as described previously ^37^. Four days later, TDLN cells were resuspended in HBSS for adoptive immunotherapy ^38,39^. Therapeutic efficacy of transferred T effector cells was assessed in the treatment of 9-day established MCA205 pulmonary metastases by intravenous injection of 5 × 10^6^ culture-activated T cells to each mouse and/or p62 plasmid (300 ug/mouse i.m) on days 9 and 14. Tumor bearing mice were pretreated intravenously with cyclophosphamide (100 mg/kg) 1 day before infusion of T cells. Cyclophosphamide treatment is routinely used to improve the therapeutic efficacy of adoptively transferred T cells and was also administered to untreated tumor-bearing control mice ^40^

### Assessment of antitumor effect

For establishment of pulmonary metastases, C57BL/6J mice were injected intravenously with either 3 × 10^5^ MCA205 suspended in 200 ul of Hanks’ balanced salt solution (HBSS). On day 26 after inoculation, MCA205 tumor–bearing lungs were counterstained with India ink and were enumerated. Lungs with more than 250 nodules were assigned >250 as the maximum number that can be counted reliably.

## Results

### Anti-tumor activity of p62 DNA plasmid in dogs & histopathological changes

The anti-tumor effects of p62 DNA has been evaluated in a cohort of six (6) dogs bearing *simple* mammary tumors (Tab. 1).

As shown in Tab. 2, the new trial confirms previous results ^24^ with p62 DNA-treated patients showing a marked reduction of the sizes of their original neoplastic masses without complete tumor eradication.

**Tab.2.** Anti-tumor activity of p62 DNA plasmid. Tumor volume for each dog was assessed 1 week after their last plasmid injection. Next, microscopic examinations revealed that the p62 plasmid induced complex changes in the histopathological patterns of original tumors. As summarized in Tab.3., two patients (# 4 and #6) demonstrated the switching of their tumor histotype from simple high malignant lesions to less aggressive complex carcinoma after p62 DNA administration. In one patient (#5), a low malignant tubulo-papillary carcinoma reverted to a benign adenoma. Patient #1 showed a transition from solid to tubulo-papillary histotype, while in patient (# 3) the residual tumor maintained the same histotype and grade as that of the original lesion. Importantly, all p62-treated patients, are still tumor- and metastasis-free and maintain good quality of life 4 years after surgery (mastectomy).

| **Pts #** | **Dose x injection (mg)** | **# injections** | **Initial** | **Final** | **Change Tumor volum. (%)** | **Mastectomy** |
| --- | --- | --- | --- | --- | --- | --- |
| **1** | **0.75** | **9** | **8,6 x 6.2** | **6,4 x 3,4** | **-78** | **YES** |
| **2** | **0.75** | **3** | **1,1 x 0,7** | **0,5 x 0,6** | **-66** | **YES** |
| **3** | **1.5** | **9** | **20 x 16** | **19 x 12** | **-75** | **YES** |
| **4** | **1.5** | **3** | **2,7 x 3,5** | **2,7 x 2,7** | **-40** | **YES** |
| **5** | **0.75** | **9** | **4,9 x 3,5** | **3 x 3** | **-55** | **NO** |
| **6** | **0.75** | **9** | **3,2 x 1,8** | **2 x 2** | **-23** | **YES** |

**Tab.3.** Changes in tumor histotype and grade following p62 DNA treatment SC: solid carcinoma; TP: tubulo-papillary carcinoma; CC: complex carcinoma; TPA: tubulo-papillary adenoma. *MS (Malignancy Score) as obtained by combining WHO Stage and grading 32: (+++), high; (++), middle; (+), low.

| **Pts #** | **Before** | **MS** | **After** | **MS \*** |
| --- | --- | --- | --- | --- |
| **1** | **SC** | **+++** | **TP** | **++** |
| **2** | **TP** | **++** | **TP** | **+** |
| **3** | **SC** | **+++** | **SC** | **+++** |
| **4** | **SC** | **+++** | **CC** | **++** |
| **5** | **TP** | **+** | **TPA** |  |
| **6** | **TP** | **+++** | **CC** | **+** |

### p62 DNA induces *alpha*-SMA and type III Collagen in Tumor Stroma

As we review in the Discussion section, increases in the *alpha*-SMA and/or Col 3 levels may mean a better prognosis. Both proteins are downregulated by chronic inflammation. Furthermore, the p62 plasmid reduced chronic inflammation in many mouse models. Thus, we hypothesized that the p62 plasmid would increase *alpha*-SMA and/or Col 3 levels. Using immunohistochemistry, we observed that p62 DNA administration resulted in significant increases in the expression of stromal *alpha*-SMA (Fig. 1) coupled with a robust synthesis and deposition of Coll 3 in the ECM, as opposed to a next to basal expression of Col 1 (Fig.2). Both Col 1 and Col 3 levels are minimal in a normal mammary gland (Fig.3).

**Fig.1.**
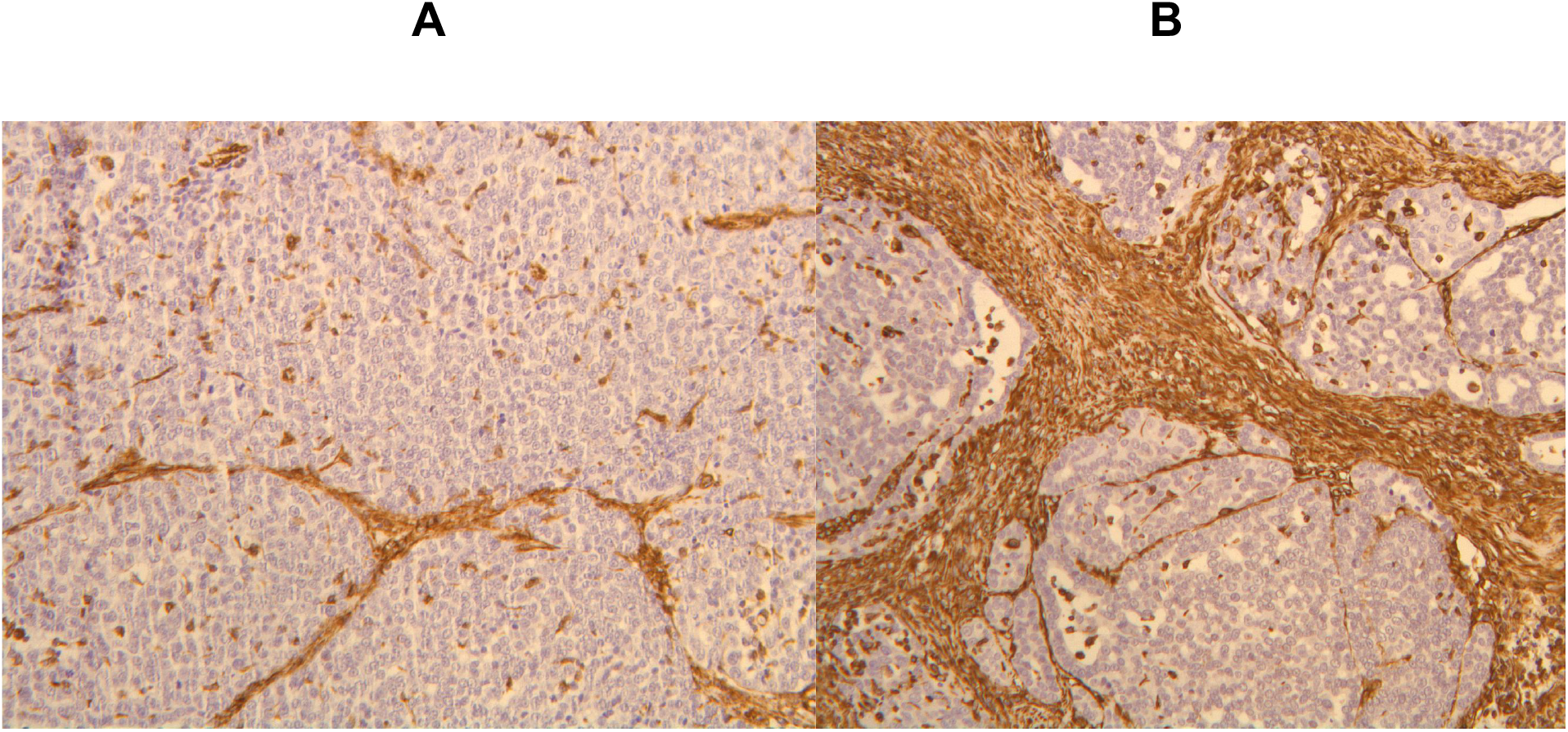
IHC staining of *alpha-*SMA in the tumor stroma before (A) and after (B) p62 DNA treatment. (20 x)

**Fig.2.**
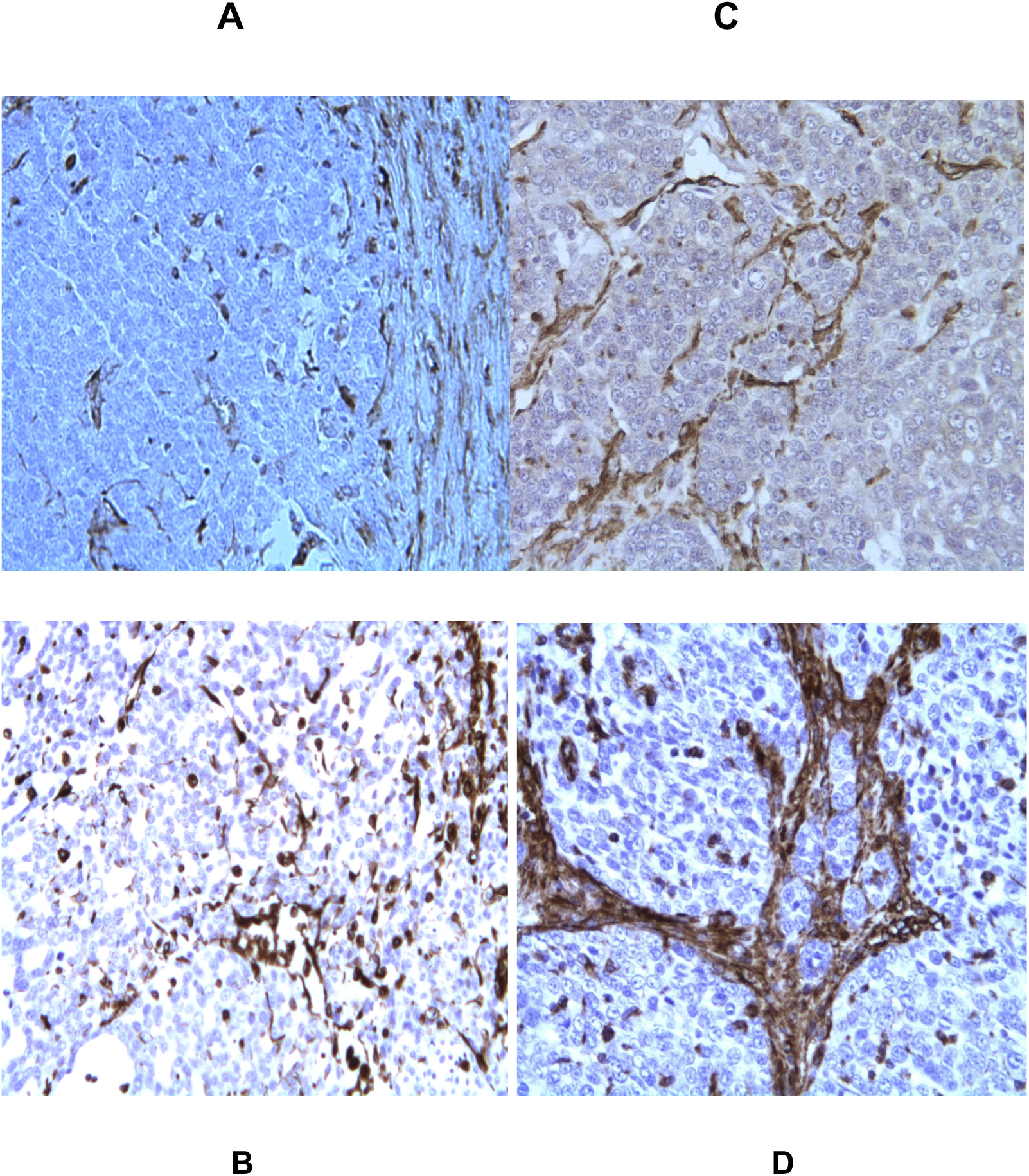
IHC evaluation of Col 1 (A, C) and Col 3 expression (B, D) in tumor biopsies, before (A,B) and after (C,D) p62 DNA injections (20x)

**Fig.3.**
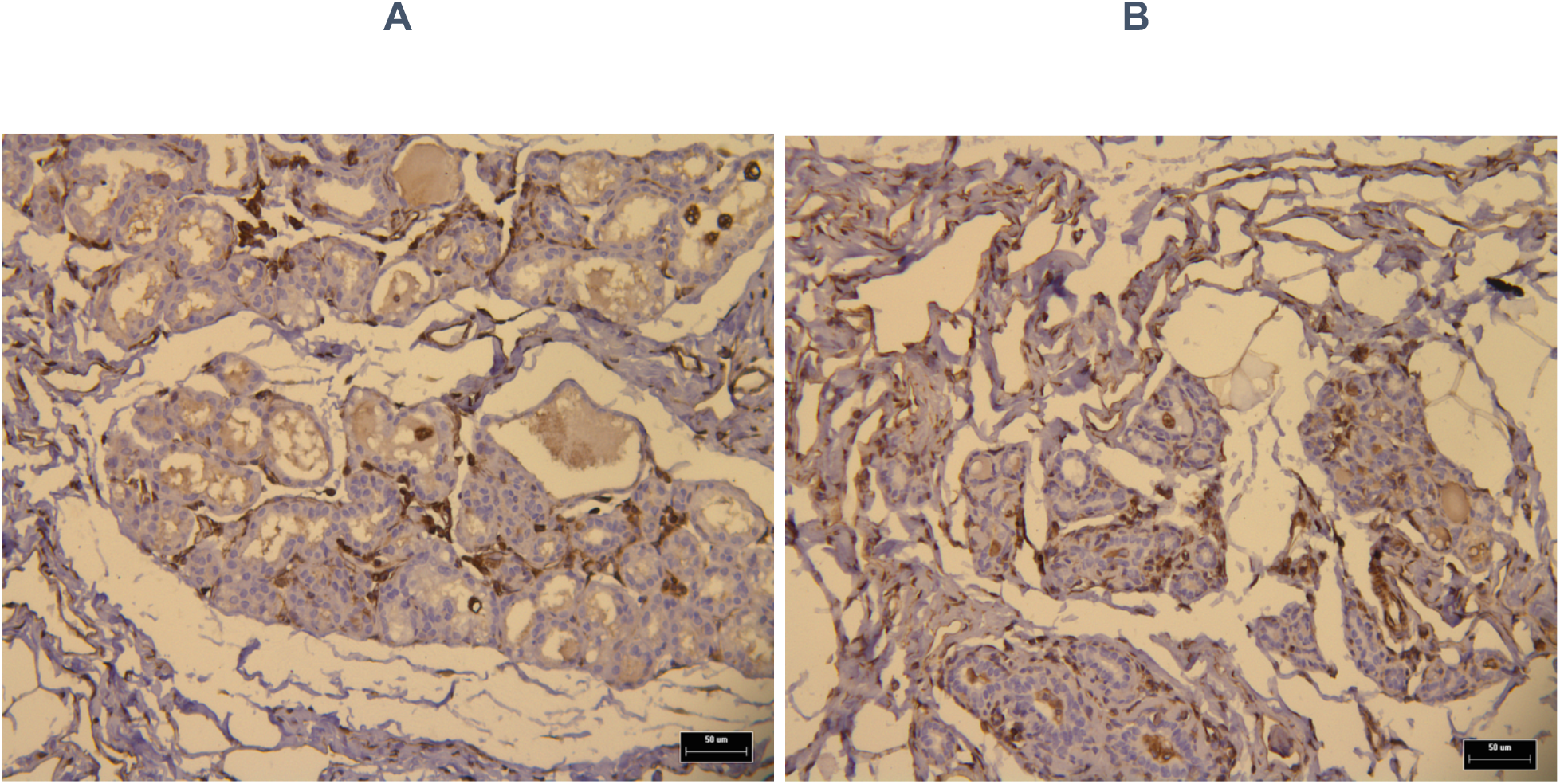
IHC of normal a mammary gland (NMG) showing both Col1 (A) and Col 3 (B) basal expression levels (brown dots)

### An adaptive immune system is indispensable for the anti-cancer effect of p62 plasmid

In dogs we have observed that p62 treatment increases the number of TILs. However, canine patients with spontaneous tumors cannot be used to learn if the increased TIL abundance is a mechanism of anti-tumor effect of p62 DNA or if it is just a coincidence. To test whether the changes in the tumor stroma induced by the p62-plasmid are sufficient for its anti-tumor effect or if the plasmid primary acts through an adaptive immunity, we compared protective effects of the plasmid in wild-type and severe combined immunodeficiency (SCID) mice. SCID mice have a genetically inactivated adaptive immune system (i.e. lacking T-and B-cells), but maintain an intact innate immune system (i.e., macrophages, NK cells etc). Thus, by comparing the antitumor effect in wt vs. SCID mouse strains one can see if an adaptive immune system is necessary for the anti-tumor effect.

When we challenged SCID mice with B16 melanoma they developed tumors similar to control (wt) syngeneic mice, indicating that the lack of an adaptive immune system does not promote tumor development in this model (Fig.4A). However, in contrast to wt mice, the p62 plasmid lost its ability to inhibit tumor growth in SCID mice (Fig. 4A). Furthermore, whereas the p62 plasmid increased survival in wt mice, no such effect was seen in SCID mice (Fig.4B). Thus, we conclude that an adaptive immune system is required for the anti-cancer effect of the p62 plasmid.

**Fig. 4A.**
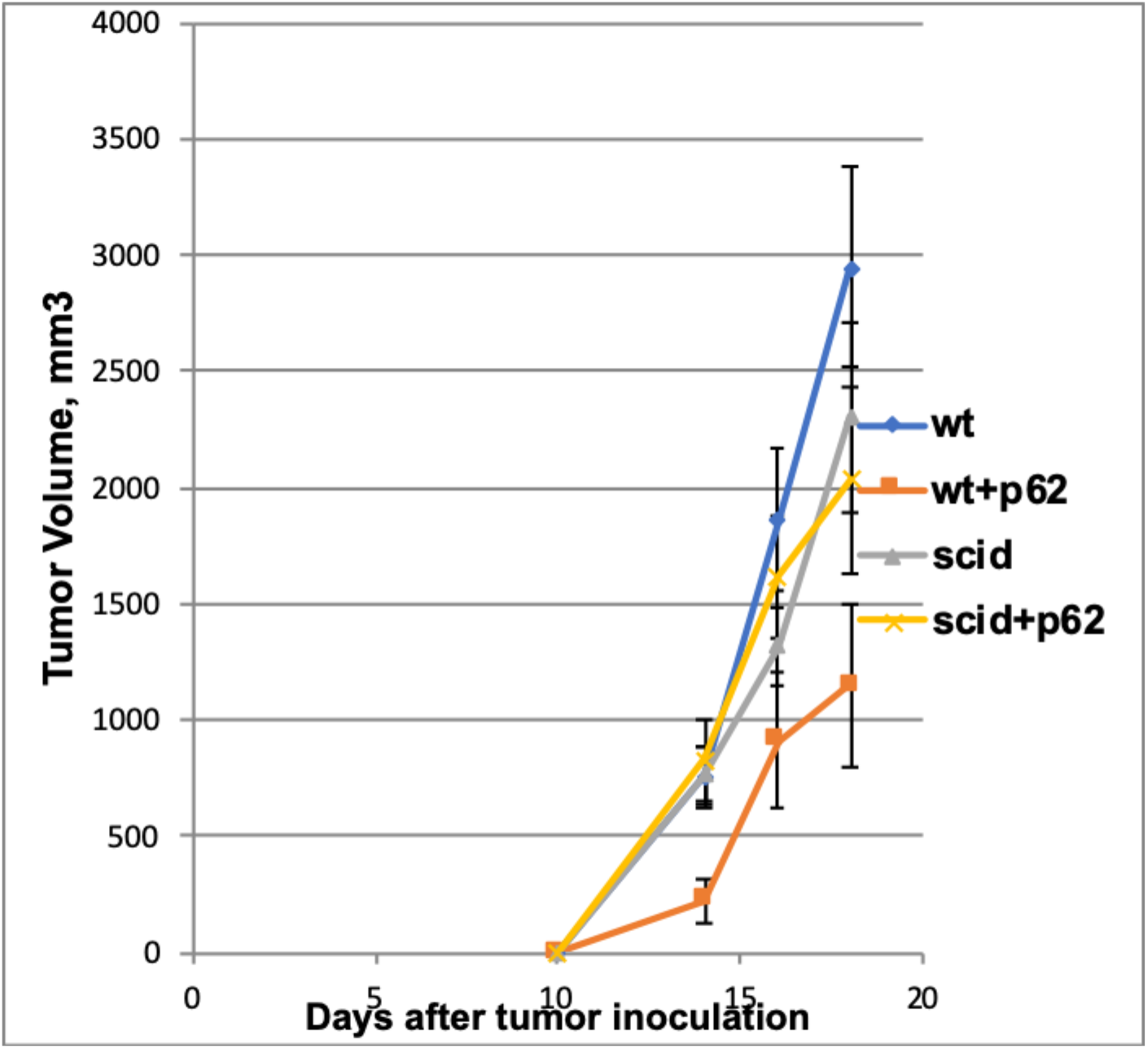
Effect of p62 plasmid on the growth of B16 melanoma in wt and immunodeficient (scid) mice. While the p62 plasmid inhibited tumor growth in wt mice, no effect was found in immunodeficient mice. wt+p62 vs wt: day14 - p= 0.008; day16 – p=0.02; day18- p=0.01 scid+p62 vs scid: day 14 - p=0.40; day 16 – p=0.31; day18 – p=0.37

**Fig. 4B.**
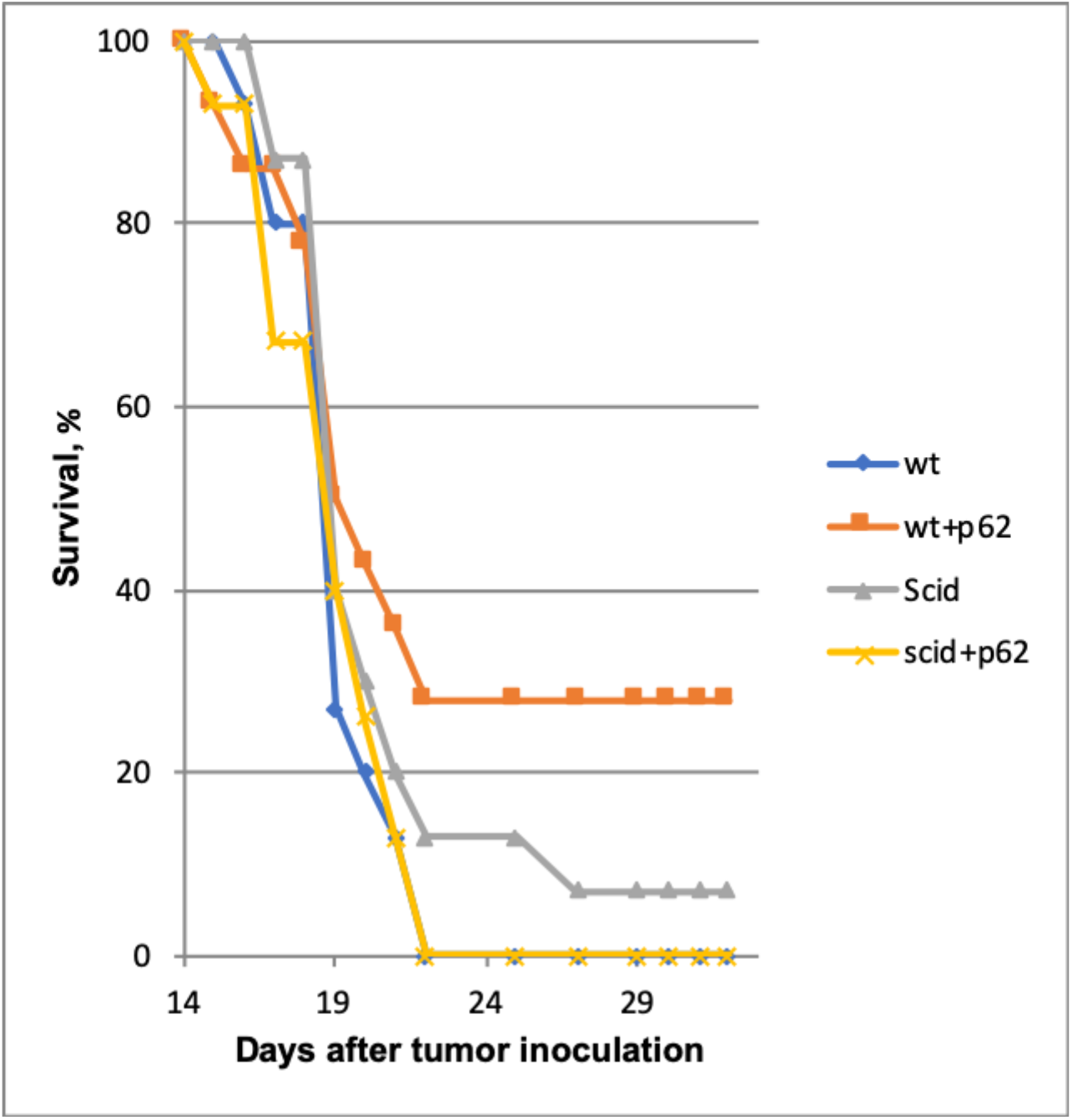
Effect of p62 on the survival of wt and immunodeficient (scid) mice. wt+p62 vs wt – p=0.026 at day 25

Thus, we established that an adaptive immune system is indispensable for the antitumor effect of p62 plasmid. Also, we knew that eliciting a specific anti-p62 adaptive immune response cannot be the only mechanism which the p62-encoding plasmid is using to stimulate anti-cancer adaptive immunity, because the plasmid is active in dogs while canine cancer cells lack their p62 (in contrast to human cells). To bring these two facts together, we hypothesized that the p62 DNA enhances effects of T-cells targeting cancer antigens other than p62 (e.g., acting via immunomodulatory mechanism). If that is correct, the p62 plasmid would increase the efficiency of any therapy generating T-cells targeting tumor associated antigens, e.g. immunotherapies. To test if the p62 DNA vaccine can enhance effects of other immune therapies, we employed a model of adaptive cell transfer where T-cells from tumor-bearing animals are transferred to animals with established tumors (or metastasis). In this model, the p62 plasmid was administered on days 9 and 14 after the animals received the transplantable tumor. This was so it did not have enough time to develop a strong protective anti-p62 immune response which could block or reduce the formation of lung metastasis. Thus, p62 DNA alone shows only a minor effect on lung metastases (Fig. 5A). At the same time, it amplified the effect of adoptively transferred T-cells on lung metastases (Fig.5B).

**Fig.5A.**
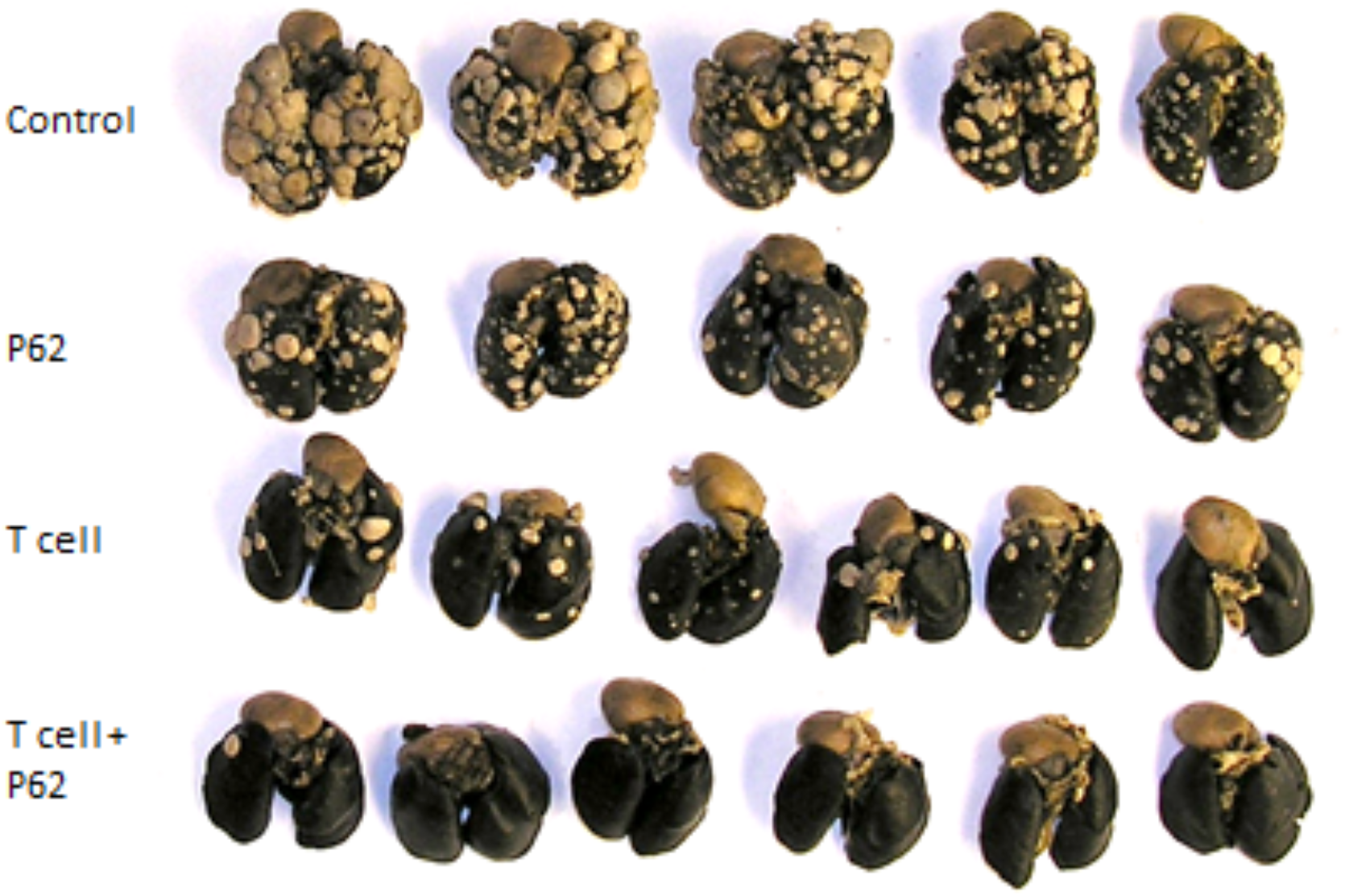

**Fig.5B.**
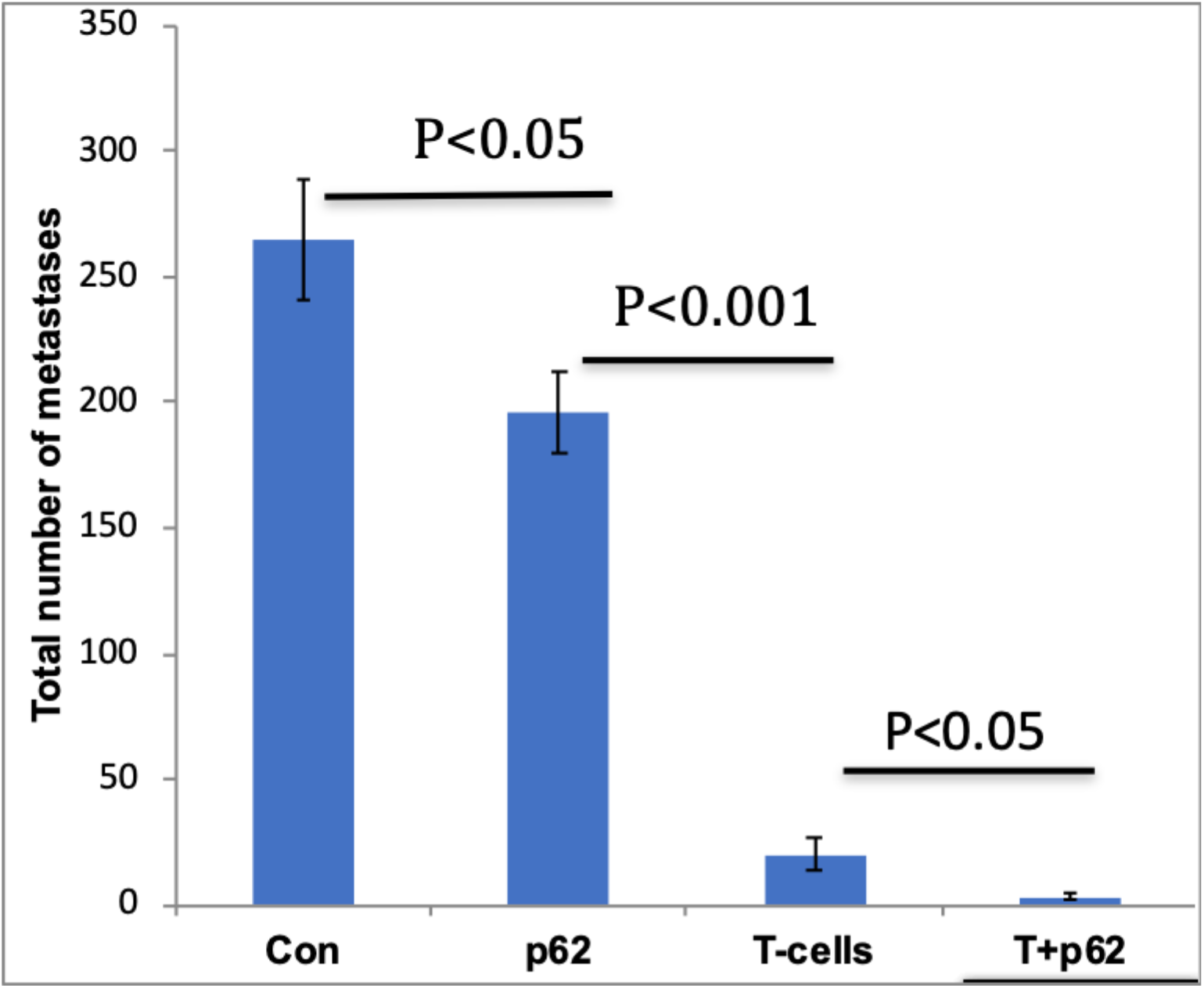
Effect of p62 plasmid on adoptive T-cell transfer. Upper panel – lung metastases formed 26 days after i.v. injection of MCA205 fibrosarcoma cells; lower panel – quantification of results.

## Discussion

The present translational oncological study was stimulated by the observations made during phase I / IIa clinical trials of patients with advanced ovarian and breast cancer. We reported transient progression free survival in a majority of patients, and a partially restored sensitivity to chemotherapy in all patients treated with the p62 plasmid ^41^. Although the causes behind the development of drug resistance include different mechanisms ^42^, growing evidence indicates that changes in tumor microenvironment may contribute to resistance against both chemotherapy and radiotherapy. In particular, importance of stroma-mediated chemosensitivity has been recognized and is the basis for the development of new anticancer agents ^43^. It is a commonly accepted view that the lower the grade of a tumor is, the more sensitive the tumor is to therapy. The results of the present paper demonstrate that the treatment with the p62 DNA induces dramatic histopathological changes in malignant tumors reverting a tumor grade towards less aggressive lesions. If the same phenomenon takes place in humans, it would explain why the p62-treated patients became responsive to chemotherapy.

A tumor stroma can generate either a tumor-permissive environment or it can constrain tumor growth by building up a “reactive” stromal phenotype characterized by alfa-Sma and Coll3 accumulation. Both Collagen 1 and 3 were reported to be downregulated under inflammatory conditions ^44^. Because p62 DNA reduces chronic inflammation (see above), we hypothesized that the anti-inflammatory properties of p62 DNA may lead to increased intratumoral expression of both proteins. Indeed, Collagen 3 expression levels were greatly increased in tumor ECM of p62 DNA treated dogs. At the same time, the fact that Col 1 was not upregulated to the same extent may not be too surprising. For example, matrix metalloproteinase cleave col I while col III level increases ^45^. Similarly, it was reported that TNF-alfa, a major pro-inflammatory cytokine, downregulates stromal alfa-SMA ^45^. Previously we demonstrated that p62 DNA quenches an ovariectomy-induced increase of TNF-alfa levels ^29^. Thus, we predicted that administering the plasmid would increase the *alfa*-SMA expression in tumor stroma. Our observations have supported this hypothesis.

Remodeling of the intratumoral ECM may partially explain the previously reported phenomenon of treatment with the p62 plasmid leading to an increased number of T-lymphocytes in canine tumors ^24^. Here, we demonstrate an increase in tumor sensitivity to T-cell therapy. It was suggested before that the trafficking and motility of cytotoxic T lymphocytes is guided by collagen fibers ^47^. Thus, by creating a network of EMC fibers throughout the tumor, p62-treatment may create the axis of collagen alignment, which CD8+ T cells can move along. Recently, a novel hypothesis was stipulated that ECM composition may define collective cooperated lymphocyte motility as opposed to an individual trafficking of the cells ^48^. Although this hypothesis was suggested for B-lymphocytes only, it would be interesting to test if the same takes place for T-lymphocytes and whether the p62 plasmid induces such collective lymphocyte tumor penetration.

Despite all the facts linking the extracellular matrix to TILs, it remained to be possible that treatment with the p62 plasmid acts on cancer cells through a mechanism not involving an adaptive immune response. For example, it was reported ^13^ that reduced Col3 level in heterozygous mice led to increased tumor formation *in vivo* when the mice were challenged with transplantable breast cancer model. *In vitro* data from the same paper suggested that the metastatic process is significantly increased when col3 level is reduced. The later phenomenon could not be due to TLC engagement because the *in vitro* system did not contain lymphocytes.

To establish whether the p62 plasmid indeed acts via an anti-tumor adaptive immune response, we compared the anti-tumor effect of the p62 plasmid in wt and SCID mice strains. Indeed, if the plasmid acts on the cancer cells directly and/or via a mechanism other than an adaptive immune response then the B16 melanoma administered to the two strains would demonstrate the same sensitivity to the plasmid. In contrast, if the plasmid acts via stimulating/modulating an adaptive immune response, it would be inactive in SCID mice lacking the lymphocytes. This later turned out to be the case. Although p62 DNA has reduced the growth of the subcutaneous B16 melanoma tumor and increased the rate of survival in the wt animals, the plasmid was completely inactive in SCID mice. Thus, we conclude that p62 DNA acts via adaptive immune system.

Considering the fact that, contrary to human tumors, canine breast cancers do not express p62 ^25^, the plasmid could not act as a classic DNA vaccine encoding p62 as a target tumor-specific antigen. Thus, we hypothesized that the p62 DNA enhances the adaptive immune response to tumor antigens other than p62. To test this hypothesis, we conducted an adoptive cell transfer experiment. T-lymphocytes were isolated from mice challenged with transplantable tumor models and, propagated *ex vivo*, were administered to animals bearing the same tumors. This led to a significant but incomplete reduction of the number of tumor lesions in the lungs. The murine cancer models are not p62 negative. However, we administered p62 DNA at late time points, so the animals did not have enough time to develop a protective anti-p62 immune response which could influence the formation of tumor lesions in lungs. According to our experience, this type of antigenic affect would take at least 3 weeks. Nevertheless, we observed that treatment with the p62 plasmid enhanced effects of adaptive T-cell transfer.

The fact that the p62 plasmid acts via an adaptive immune response corresponds well to the observations that the plasmid restores sensitivity to chemotherapy. Despite the fact that originally chemotherapeutic agents were selected based their ability to kill rapidly dividing cells, it turned out that many of them act via the immune system (e.g. stimulating immune-presenting cell death or regulating T-regs) ^49, 50^. Thus, the result of chemotherapy is a lymphocyte attack on cancer cells. Creating a tumor microenvironment favorable for active TILs makes p62 DNA an equally promising adjuvant for immune-, chemo- and radiation therapies because all of them involve immune response.

### Conclusions

We conclude that administering the p62-encoding plasmid changes intratumoral microenvironment and reverts tumor grade. The plasmid can be used as an adjuvant for cancer therapies directly and/or indirectly acting through an immune response (such as chemo- and immunotherapies).

## Acknowledgements

The authors would like to thank prof. Michael Sherman for intellectually stimulating discussions and editorial comments.

